# Modeling of RNA-seq fragment sequence bias reduces systematic errors in transcript abundance estimation

**DOI:** 10.1101/025767

**Authors:** Michael I. Love, John B. Hogenesch, Rafael A. Irizarry

## Abstract

RNA-seq technology is widely used in biomedical and basic science research. These studies rely on complex computational methods that quantify expression levels for observed transcripts. We find that current computational methods can lead to hundreds of false positive results related to alternative isoform usage. This flaw in the current methodology stems from a lack of modeling sample-specific bias that leads to drops in coverage and is related to sequence features like fragment GC content and GC stretches. By incorporating features that explain this bias into transcript expression models, we greatly increase the specificity of transcript expression estimates, with more than a four-fold reduction in the number of false positives for reported changes in expression. We introduce *alpine*, a method for estimation of bias-corrected transcript abundance. The method is available as a Bioconductor package that includes data visualization tools useful for bias discovery.

## INTRODUCTION

RNA sequencing (RNA-seq) provides a rich picture of the transcriptional activity of cells, allowing researchers to detect and quantify the expression of different isoforms and types of genes. To estimate transcript abundance, software such as *Cufflinks* [1] or *RSEM* [2] is used to take RNA-seq reads and (optionally) transcript annotations to infer i) which transcripts are expressed and ii) at what level. The results of downstream analyses are heavily dependent on these software and their underlying models and algorithms. Accurate and robust transcript abundance estimates are critical and enable further analyses and downstream validation.

Consortia have investigated the accuracy of RNA-seq expression estimates at scale and have confirmed that significant batch-specific biases are often present, that relative expression across samples is more reproducible than absolute expression estimates, and that batch effects arise mostly from RNA extraction and library preparation steps [3-5]. Computational methods for estimating gene and transcript abundance attempt to mitigate the effect of technical biases by estimating sample-specific bias parameters with the hopes of capturing as much unwanted variation as possible. For gene-level expression, common normalization methods include fitting a smooth curve to the observed counts of RNA-seq fragments for each gene over the GC content and length of the genes [6, 7]. Other methods target batch effects directly at the gene-level counts using approaches based on factor analysis [8-10]. However, here we are interested in transcript-level analysis not the gene-level analysis these methods address.

At the transcript level, a number of sample-specific bias parameters are estimated by methods and included in the model during the estimation of transcript abundance. These known biases and their sources include the fragment length distribution from size selection, the positional bias along the transcript due to RNA degradation and mRNA selection techniques, and a sequence-based bias in read start positions arising from the differential binding efficiency of random hexamer primers [2, 11-15] (Figure 1a). Some methods apply a correction to transcript-level estimates after abundance estimation using the average GC content of the transcripts [16, 17].

**Figure 1:**
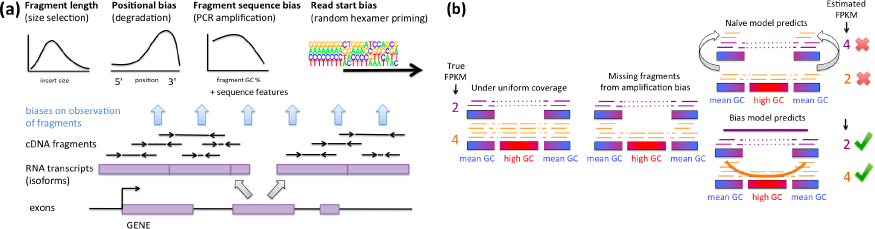
(a) RNA-seq biases. (b) Ignoring fragment sequence bias impairs transcript abundance estimation when critical regions that distinguish isoforms have GC content or sequence features that make fragments hard to amplify, resulting in false positives of predicted expression of isoforms that are lowly or not expressed.

Recent investigations into the coverage of RNA-seq fragments along transcripts revealed examples of extreme variability in coverage that is purely technical and sample-specific [18]. Despite these efforts toward transcript-level bias correction, highly variable patterns of coverage confound current methods that are designed to identify and quantify transcripts [19].

Here we investigate the cause of systematic errors in transcript abundance estimates. We find that GC content and other sequence features in the fragment explain much of the coverage variation within transcripts, and outperform the read start features used by current methods to reduce the bias from random hexamer priming. Despite the importance of GC content in explaining differences in expression estimates across batches, none of the existing transcript abundance estimation methods incorporate sample-specific fragment-level GC content bias in their models [3, 5]. We show that ignoring these fragment sequence biases can lead to hundreds of false positive estimates of transcript expression and misidentification of the major isoform, as methods are unable to interpret drops in coverage as technical artifacts (Figure 1b). By incorporating a term for fragment sequence bias alongside other bias terms into an expectation-maximization algorithm for transcript abundance estimation, we are able to produce estimates that are more stable across centers and batches. The bias modeling framework and transcript abundance estimation methods are distributed as an open-source R/Bioconductor package *alpine.* Our framework enables further research both into optimization of library preparation protocols to reduce or eliminate biases as well as computational approaches that mitigate bias.

## RESULTS

### Current methods predict many significant differences across center

We sought to identify and quantify different kinds of technical bias in RNA-seq data, both on absolute estimates of abundance and in comparisons of expression estimates across samples. To uncover such biases, we downloaded 30 RNA-seq samples from the GEUVADIS Project of lymphoblastoid cell lines derived from the TSI population, 15 of which were sequenced at one center and 15 at another [20] (Supplementary Table 1). We ran state-of-the-art transcript quantification software *Cufflinks* (with the sequence bias removal option turned on) and *RSEM* on these 30 samples, generating a table of estimates for RefSeq transcripts. Performing a simple t-test on log_2_(FPKM + 1) values from *Cufflinks* across centers, and filtering on Benjamini-Hochberg adjusted *p* values less than 1% false discovery rate (FDR) resulted in 2,510 transcripts out of 25,588 with FPKM greater than 0.1 reported as differentially expressed (Figure 2a, see Supplementary Figure 1 for *RSEM* results). 619 out of 6,761 genes with multiple isoforms and FPKM greater than 0.1 for one or more isoform had changes in the reported major isoform across centers using FPKM values estimated by *Cufflinks* (Supplemental Table 2). While it is expected that there will be batch effects when comparing estimates from samples prepared in different centers, our across-center analysis provides a baseline picture of the extent of systematic errors in absolute transcript-level abundance estimates. Well-designed sequencing projects such as GEUVADIS are careful to distribute samples from one biological condition (here population) across centers and to consider this in statistical comparisons.

**Figure 2.**
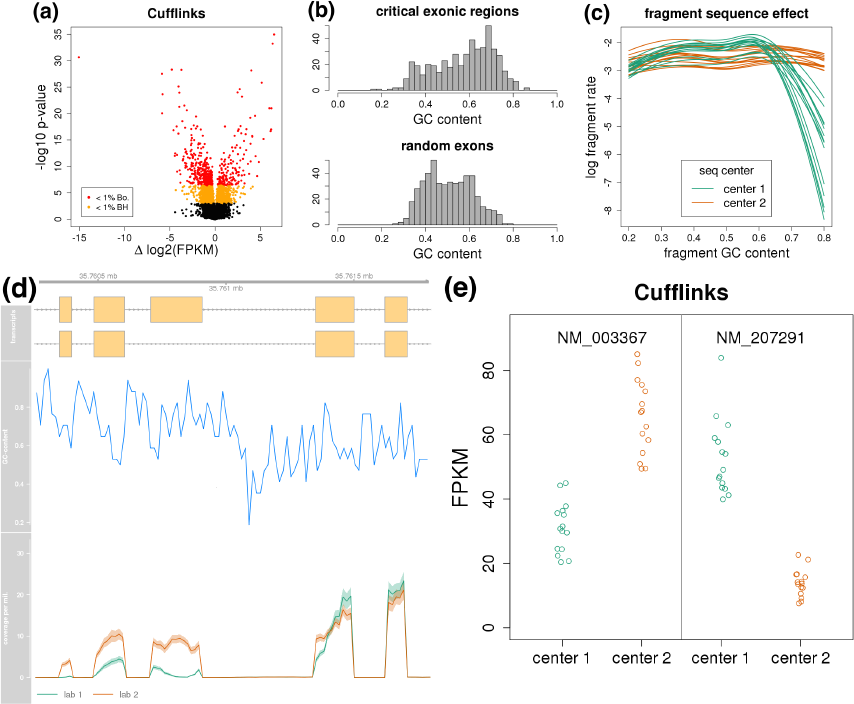
Problems with current transcript abundance estimation methods. (a) Volcano plot of a comparison of *Cufflinks* transcript estimates across center, with 2,510 transcripts having FDR less than 1% and 515 with family-wise error rate (FWER) of 1% using a more conservative Bonferroni correction. (b) GC content of critical gene regions distinguishing two isoforms when one or more reported differential expression, compared to GC content of random exons. (c) Dependence of fragment rate on GC content after controlling for random hexamer priming bias of read starts. (d) Coverage and GC content in a region of the USF2 gene containing the alternative exon. (e) *Cufflinks* FPKM estimates for USF2.

We then examined what sources of technical bias might be underlying the differences in transcript abundance estimates. While the transcripts with reported differential expression were equally divided among single and multiple isoform genes as the rest of transcripts, we choose to focus on genes with two isoforms, so that we could more easily identify what features of genes might be causing the difference in estimated expression. Out of 5,716 transcripts from genes with two isoforms in which at least one isoform had FPKM greater than 0.1, 566 transcripts reported differential expression across center according to *Cufflinks* estimated abundances, at an FDR threshold of 1%. Of these, 164 transcripts were from genes where both of the two isoforms were reported differentially expressed, and in the majority of cases the reported fold change was in different directions (134 different vs 30 same direction). Furthermore, the critical regions of the genes’ exonic structure - those regions that were exclusive to one or the other isoform - for genes with one or more isoform reported as differentially expressed, had much higher GC-content (mode at 70% compared to 50%) than expected by chance (Figure 2b, Wilcoxon *p* < 0.0001).

An example of a gene with differences in expression estimates across center is USF2 (Figure 2d). It is often small, critical regions (exons or parts of exons) that distinguish isoforms of a gene. For some genes, these regions will have stretches of high GC content. Because methods such as *Cufflinks* and *RSEM* employ a likelihood model that ignores differences in coverage due to fragment sequence features like GC content, the drop in coverage for samples from center 1 results in a shift in expression estimates from the first isoform to the second isoform, which does not include the high GC exon (Figure 2e). Note that the samples from center 1 have dramatically reduced representation of high GC fragments compared to center 2, even after adjusting for differences due to random hexamer priming bias (Figure 2c).

### A model for RNA-seq fragment sequence bias

PCR amplification of DNA fragments generates the GC content bias seen in sequencing data [21-23], and for DNA-seq it has been shown that optimal correction for the amplification bias occurs when modeling at the scale of the fragment [24]. Therefore, we used fragment features, such as fragment GC content and the presence of long GC stretches within the fragment (defined in Methods), to predict the number of times a potential fragment of a transcript was observed (0,1, 2,…). The effect of GC stretches within a gene in reducing RNA-seq coverage has been previously described [25]. We considered all potential fragments with a range of length within the center of the fragment length distribution, at all possible positions consistent with the transcript’s beginning and end. While one existing method for estimating percent of isoform expression assumed a positive linear relationship between exon counts and exon GC content [26], our investigation reconfirmed the findings of Benjamini and Speed [24], that the GC content effect was often nonlinear and highly sample-specific, therefore requiring sample-specific estimation of smooth curves of GC content.

We constructed a Poisson generalized linear model for the count of each potential fragment for each sample separately, with a number of modular bias correction terms that could be used exclusively or in combination (see Methods for details). Possible terms included smooth curves for the fragment GC content, indicator variables for the presence of GC stretches, a term for the fragment length, smooth functions of the relative position within the transcript, and the read start bias for both ends of the fragment, using the same variable length Markov model (VLMM) proposed by Roberts et al. [13] and used in *Cufflinks.*

### Fragment sequence explains more technical variability in coverage than read start sequence

To test the predictive power of various models for technical bias, we downloaded a recently published benchmarking dataset, where 1,062 human cDNA clones were *in vitro* transcribed and the resulting transcripts were mixed at various concentrations with mouse total RNA, prepared as libraries and sequenced (IVT-seq) [18]. Libraries were prepared using standard techniques, including poly(A) enrichment, no selection, and for those libraries titrated with mouse total RNA, ribosomal RNA depletion was performed. The resulting publicly available dataset is valuable for studying technical variability in coverage, as the exact sequence of the transcripts are known, and the raw data displays the typical non-uniform transcript coverage not found in spiked-in DNA. We focused our analysis on 64 of the IVT transcripts defined by Lahens et al. [18] as exhibiting “high unpredictable coverage”.

We compared the predictive power of various models, all of which included fragment length, and which optionally included read start sequence bias, fragment GC content, and identifiers for GC stretches (Supplementary Figures 2-4). For prediction, we used 2-fold cross validation, such that two models were trained on two halves of the data, and always evaluated on transcripts that were not in the training set. Predictive power was measured as the percent reduction of mean squared error in explaining raw fragment coverage, compared to a null model that predicted uniform coverage across the transcript. The models that included fragment GC content doubled the predictive power of the model that included read start sequence bias (Figure 3a). The model that also included the information about GC stretches was more predictive than the model with just the fragment GC content, although only slightly so. The fragment sequence models accurately capture the drops in coverage that were not captured by the read start sequence model (Figure 3b, Supplementary Figures 5-6).

**Figure 3.**
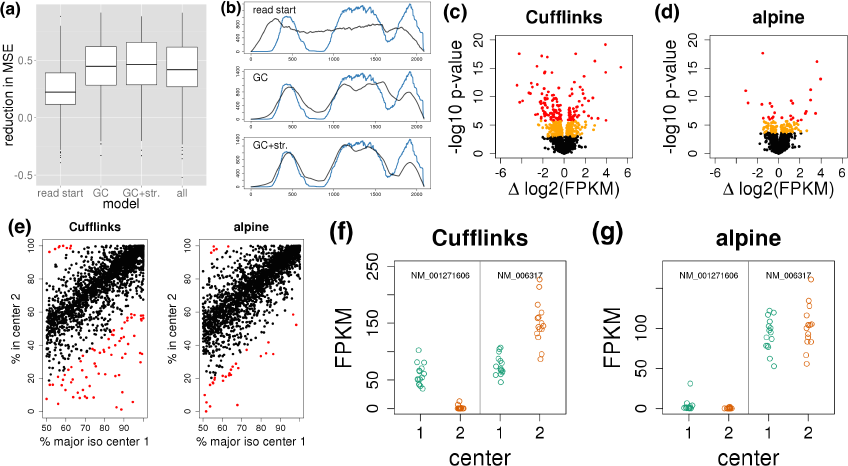
Modeling and correcting fragment sequence bias. (a) Comparison of reduction in mean squared error (MSE) for different bias models for all 8 samples and 64 transcripts (n=512). (b) Comparison of test set coverage prediction for bias models on GenBank BC011380 (raw coverage in blue, predicted coverage in black). (c) and (d) Volcano plots of differential transcript expression across centers for genes with two isoforms (orange: Benjamini-Hochberg FDR less than 1%; red: Bonferroni FWER rate less than 1%). (e) Consistency of percent expression of the major isoform identified in center 1 compared to center 2. *Cufflinks* had 78 genes with estimated isoform percent change more than 35% across center, while *alpine* had 32 genes (red points). (f) and (g) FPKM estimates for two isoforms of BASP1 across center.

### Correcting for fragment sequence bias reduces false positives of estimated transcript expression

We then used our approach to compare the transcript abundance estimates from one center against the other in the 30 GEUVADIS samples. To more clearly show the performance with respect to differential isoform usage, we focused on 5,712 transcripts from genes with only two isoforms and FPKM values estimated by *Cufflinks* greater than 0.1 for one of the two isoforms. Additionally, we required that the two isoforms have at least one overlapping basepair. We compared log_2_(FPKM + 1) estimates across center using a t-test. We found that including bias terms for fragment GC and GC stretches resulted in more than a four fold decrease in the number of false positives at an FDR threshold less than 0.01 *(Cufflinks* reported 562 differentially expressed transcripts, while *alpine* reported 130 (Figure 3c-d)). Using a more conservative Bonferroni correction, *Cufflinks* reported 157 transcripts differentially expressed transcripts across center with FWER of 1%, while *alpine* reported only 27. Of the 5,712 transcripts considered, *Cufflinks* reported 5,020 with FPKM greater than 0.1 and *alpine* reported 4,903. In general, *alpine* greatly reduced across-center significant differences while within-center coefficient of variation of abundance estimates remained the same as for *Cufflinks* (Supplementary Figures 7-10).

Likewise we observed reduced across-center differences for estimation of isoform percentages within the 2,856 genes. For each gene, we calculated the estimated percent expression of the major isoform for center 1 (a number ranging from 50% to 100% by definition), against the estimated percent expression of that same isoform in center 2. Comparing *Cufflinks* with *alpine*, inclusion of the fragment sequence terms reduced the number of extreme predicted changes in isoform percent when comparing across center (Figure 3e). An example of how false positives for isoform switching arose is the two-isoform gene BASP1, with the FPKM estimates from both methods shown in Figure 3f-g (additional examples in Supplementary Figure 11). Similar improvements were attained over *RSEM* (Supplementary Figures 12-13). Including read start bias terms in the *alpine* model did not provide visible improvements (Supplementary Figures 14-17).

Fitting the fragment sequence model does require more computational effort than models which assume uniformity of fragments, or which correct only for fragment length distribution, though our implementation runs in comparable time to the *cuffquant* and *cuffnorm* steps of the *Cufflinks* suite. Generating the bias coefficients for the 30 GEUVADIS samples required 24 minutes using 6 cores and 75 Gb of memory. Estimating the transcript abundances for 5,712 transcripts from two-isoform genes for all 30 samples required 3.5 hours using 40 cores and 50 Gb of memory. The *alpine* software is implemented using core Bioconductor packages and the *speedglm* R package [27, 28].

## DISCUSSION

Systematic errors and batch effects are a continuing cause of concern for RNA-seq experiments. Large-scale, well-documented transcriptome sequencing projects such as GEUVADIS [20] allow the creation of computational methods that correct technical biases, such as sample-specific fragment sequence bias. Here we used an across-center comparison to find and quantify significant differences in transcript expression estimates. We note that our findings reflect general systematic errors and not just differences induced by batch effects. The problem holds with absolute abundance estimates derived from data from a single center. There are likely to be many incorrectly reported major isoforms and biased abundance estimates for experiments that show strong dependence of the fragment rate on GC content (e.g. Figure 2d), unless these are explicitly corrected for using fragment sequence bias modeling.

For analyses focusing on differential abundance across conditions, batch effects affecting a number of transcripts can be controlled for with proper experimental design - distribution of samples from one biological condition across centers - and by considering technical factors in statistical comparisons. However, for those genes with misidentification of the major isoform, for example, the more than one hundred transcripts we identified from genes with two isoforms that showed significant and opposite directions of expression fold changes (Figures 2e and 3f), including blocking or inferred technical variation factors will not prevent a differential signal across condition being attributed by a naive model to the wrong transcript.

While computational methods are available for correcting gene-level expression using average GC content for the gene [6, 7] or by estimating hidden factors [8-10], the systematic errors induced at the transcript level are more problematic and difficult to correct for than at the gene level. Transcript-level corrections do not effectively correct the bias, because the GC content of the isoforms can be very similar when the critical regions distinguishing isoforms are short compared to the total transcript length. Removal of duplicate reads/fragments, and barcoding of unique molecules is also not likely to correct for these technical biases, as the observation of one fragment compared to zero is equally affected by PCR amplification bias as the observation of two or more compared to one. Longer reads or fragments are also unlikely to correct this bias, as from our investigation, PCR amplification is impaired based on sequences contained within fragments. Here we focused primarily on specificity, showing a decrease in false positives through modeling of additional bias terms. New benchmarking experiments are necessary in order to test sensitivity: experiments where the true isoform or set of isoforms are known, and in which characteristically highly-variable profiles of transcript coverage are obtained by following as closely to the steps of a standard RNA-seq experiment as possible.

While the sequence features we included in our model provided substantial improvements over existing methods, we hypothesize that more variability can be explained by discovering new predictive features. Our R/Bioconductor package provides a modular framework that facilitates further exploration. For example, though we did not include an interaction term between fragment length and GC content curves as described by Benjamini and Speed [24], this could be added. Our software also will prove useful for optimization of protocols to reduce GC content bias [22] or bias due to stretches of GC, which is preferable to computational corrections.

## METHODS

### RNA-seq read alignment

IVT-seq FASTQ files made publicly available by Lahens et al. [18] were downloaded from the Sequence Read Archive. Paired-end reads were aligned to the human reference genome contained in the Illumina iGenomes UCSC hg19 build, using STAR version 2.3.1 [29]. The exons of the GenBank transcripts were read from the 

~~~
feature_quant.txt
~~~

 files posted to GEO by Lahens et al. [18]. The list of transcripts with high unpredictable coverage was downloaded from the additional files of Lahens et al. [18].

GEUVADIS FASTQ files made publicly available by Lappalainen et al. [20] were downloaded from the European Nucleotide Archive (see Supplementary Table 1). Paired-end reads were aligned to the human reference genome contained in the Illumina iGenomes UCSC hg19 build, using TopHat version 2.0.11 [30]. The 

~~~
genes.gtf
~~~

 file contained in the Illumina iGenomes build was filtered to genes on chromosomes 1-22, X, Y and M, and provided to *Cufflinks, RSEM* and *alpine* as gene annotation.

### Transcript quantification for GEUVADIS

*Cufflinks* version 2.2.1 [1, 13] was run with bias correction turned on, with the commands:

~~~
cuffquant -p 40 -b genome -o cufflinks/file genes.gtf \
    tophat/file/accepted_hits.bam
cuffnorm genes.gtf -o cufflinks cufflinks/file1/abundances.cxb \
    cufflinks/file2/abundances.cxb …
~~~

*RSEM* version 1.2.11 [2] was run with the commands:

~~~
rsem-prepare-reference —gtf genes.gtf genome rsem/hg19
rsem-calculate-expression -p 20 --no-bam-output --paired-end \
    <(zcat fastq/file_1.fastq.gz) <(zcat fastq/file_2.fastq.gz) \
    rsem/hg19 rsem/file/file
~~~

### RNA-seq fragment sequence bias model

The following model applies to paired-end RNA-seq fragments, but could be easily modified for single-end reads. For each sample and each transcript (each isoform of a gene), we build a 2 dimensional matrix *Y* in which we store counts (0,1, 2,…) of the aligned paired-end fragments. This matrix is indexed along the rows *p* by the start of the first read of a potential fragment, and along the columns *l* by the length of a potential fragment. Fragments which were not observed will then have *Y*_*pl*_ = 0. For computational efficiency, the columns are limited to the middle 99% of the empirical fragment length distribution. For the IVT-seq samples, the center of the distribution was defined by *l* ∊ [100, 350] and for the GEUVADIS samples, *l* ∊ [80, 230], with a total of L fragment lengths considered. The elements of the matrix *Y*_*pl*_ are then the counts of fragments with a read starting a position *p* and of length l. Note that most entries of this matrix are 0. For estimating bias parameters and comparison of predicted coverage to observed coverage, the fragments which begin on the first basepair or end on the last basepair of a transcript are not included, as large counts for these potential fragments could impair estimation of the coefficients for coverage biases within the body of the transcript. The matrix *Y* represents nearly all of the potential fragment types which could occur from a given transcript. Paired-end reads which are compatible with multiple isoforms are assigned a 1 to each of the transcript matrices *Y*, as long as the fragment length is within the range defined above. The counts in the matrix are modeled on a number of features including:

- The fragment’s length
- The relative position of the fragment in the transcript
- The GC content of the fragment (and other features of the fragment sequence)
- The sequence in a 21 bp window around the starts of the two paired-end reads. This is used by the *Cufflinks* variable length Markov model (VLMM) for the random hexamer priming bias [13].

The counts in the matrix Y are collapsed into a vector 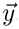, which is indexed with *j* such that *y*_*j*_ is a count for the *j*-th potential fragment type. Data for all of the features listed above is gathered for all the potential fragments using vectorized Bioconductor functions [27]. Two offsets are calculated using the observed fragments aligning to genes with a single isoform: the length of a particular fragment is converted into an offset using the log of the empirical probability density function, and a 21 bp VLMM is calculated and turned into an offset identical to the one introduced by Roberts et al. [13] for *Cufflinks.*

A number of coefficients are then included in a Poisson generalized linear model (GLM). These coefficients include: a natural cubic spline based on the GC content for each fragment (with knots at 0.4,0.5,0.6 and boundary knots at 0,1), a natural cubic spline for the relative position of the fragment in the transcript (with knots at 0.25, 0.5, 0.75 and boundary knots at 0, 1), and four indicator variables which indicate if the fragment contains a stretch of higher than 80% or 90% GC content in a 20 or 40 bp sequence within the fragment. These terms together form a model matrix *X*, where *X*_*j*_. gives the row of the model matrix for the *j*-th potential fragment. For fitting the GLM across multiple genes, an indicator variable is added for the different genes used in the bias modeling steps. This term absorbs any differences due to gene expression. Note that we do not model an interaction between fragment length and GC content of the fragment, as discussed by Benjamini and Speed [24], because we were able to predict coverage drops with GC content alone and so to limit the number of parameters in the model. However, the interaction could be added at this stage.

*y*_*j*_ is modeled as follows:

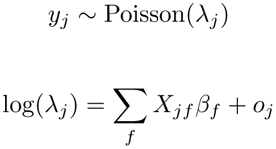

where *f* indexes the columns in the matrix *X* and **β**_*f*_ is the matching coefficient. The *O*_*j*_ term is the offset from fragment length and/or the variable length Markov model (VLMM) term for the read start primer bias from both reads. The coefficients and offsets are estimated using the genes that have only one isoform, which avoids the problem of probabilistic deconvolution of the fragments from different isoforms of a single gene. The coefficients and offsets were estimated across the 64 transcripts in the IVT-seq dataset (in two batches for cross-validation) and across 60 medium to highly expressed genes in the GEUVADIS dataset.

### Predicted read start coverage

The predicted read start coverage for position *p* is defined as:

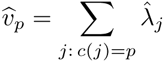

where *c(j)* = *p* indicates that the *j*-th potential fragment covers position *p*. Note that the predicted coverage 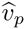 is only used for plotting and model comparison on the IVT-seq dataset, and not for estimation of the GLM coefficients or transcript abundance. For estimation of the GLM coefficients and transcript abundance, only the fragment-level counts *y*_*j*_ and estimates 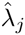 are used.

### Bias models

The following models were fit for IVT-seq samples:

- “GC”: *y* ∼ gene + frag. length + frag. GC content
- “GC+str.”: *y* ∼ gene + frag. length + frag. GC content + GC stretches
- “read start”: *y* ∼ gene + frag. length + VLMM
- “all”: *y* ∼ gene + frag. length + frag. GC content + GC stretches + VLMM

The following models were fit for GEUVADIS samples:

- “GC+str.”: *y* ∼ gene + frag. length + rel. position + frag. GC content + GC stretches
- “read start”: *y* ∼ gene + frag. length + rel. position + VLMM
- “all”: *y* ∼ gene + frag. length + rel. position + frag. GC content + GC stretches + VLMM

In the IVT-seq dataset, relative position was not included, as strong positional bias was not observed, as opposed to the poly(A)-selected GEUVADIS dataset, which did exhibit positional bias. During modeling, the observations from highly expressed genes are down-sampled, so that each gene contributes equally to the final model in terms of FPKM. This is equivalent to the approach in Roberts et al. [13] for the *Cufflinks* bias estimation steps.

### Transcript abundance estimation

The estimated bias terms can be used to improve the estimates of transcript abundance for genes with single isoforms or multiple isoforms. For fragment type *j* and isoform *i*, the Poisson rate is given by:

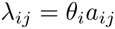

where *A* is a sampling rate matrix, and 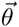 represents the abundance of the different isoforms of a gene, as described by Salzman et al. [31] and Jiang and Salzman [32]. *a*_*ij*_ = 0 if fragment type *j* could not arise from isoform *i*. If fragment type *j* can arise from isoform *i*, one parametrization sets *a*_*ij*_ = *q(l*_*j*_)*N*, where *q* is the empirical density of fragment lengths, *l*_*j*_ is the fragment length of the *j*-th fragment type, and *N* is the total number of mapped reads. Here, we included the fragment length in the overall bias term 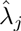, so we set 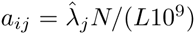 when fragment type *j* can arise from isoform *i*. The denominator contains 10^9^ and the range of the fragment lengths *L* considered in the model, so that final estimates of 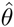 are on the FPKM scale.

Note that 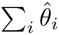 is not equal to 1, as these represent expression abundances, so they are only required to be non-negative (the 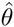 are proportional to FPKM). The Poisson model defined here is the same model as proposed by Jiang and Salzman [32], but here we explicitly model the bias using offsets 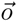, a matrix of features *X*, and a vector of log fold changes 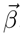, so not the same *β* as defined by Jiang and Salzman [32].

The log likelihood of a given estimate of the isoform abundances is evaluated with:

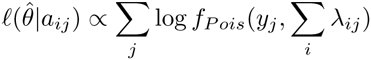

The maximum likelihood estimate of 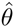 is obtained using an EM algorithm as described by Jiang and Salzman [32], where fragment types *j* with no observed fragments need only be considered for certain steps of the EM. For genes with a single isoform, the maximum likelihood estimate for *θ*, given the estimated bias terms *λ*_*j*_, is:

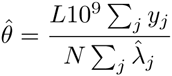

As some of the bias terms introduce arbitrary intercepts, the estimates 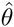 for all transcripts are scaled by a single scaling factor for each sample to match the null model estimates (the model with *λ*_*j*_ = 1) using the median ratio of 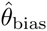 over 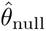. For normalizing transcript abundances across samples, the 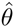 estimates are scaled using the median-ratio method of *DESeq* [33].

## AVAILABILITY OF SOFTWARE

The *alpine* software is available at the following repository: 

~~~
https://github.com/mikelove/alpine
~~~

.

## ACKNOWLEDGMENTS

The authors are grateful for helpful suggestions from Yuval Benjamini, Wolfgang Huber, Nicholas Lahens, Luca Pinello and Clifford Meyer. MIL is supported by NIH grant 5T32CA009337-35. JBH is supported by NIH R01 grant HG005220, the National Institute of Neurological Disorders and Stroke (5R01NS054794-08 to JBH), the Defense Advanced Research Projects Agency (DARPA-D12AP00025, to John Harer, Duke University). RAI is supported by NIH R01 grant HG005220.

## Supplementary Tables and Figures

### Supplementary Tables

**Supplementary Table 1:**
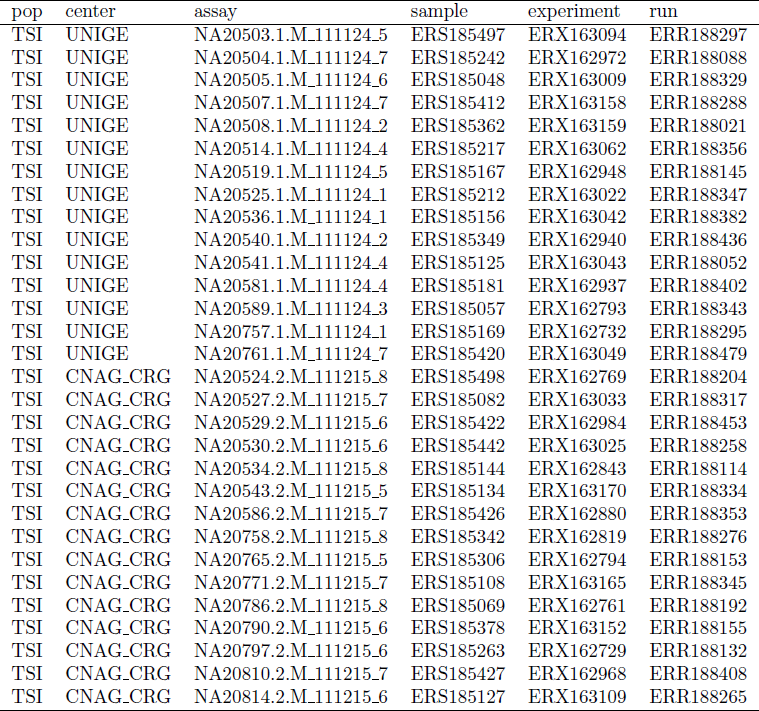
Information on the GEUVADIS samples. CNAG_CRG was coded as center 1 and UNIGE was coded as center 2 in the text. The read length was 75 for all samples.

**Supplementary Table 2:**
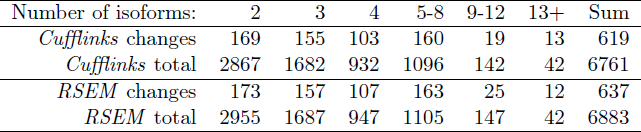
Number of genes with changes in major isoform. Considering genes which have more than one isoform, and which had estimated FPKM greater than 0.1 in at least one isoform (total), shown is the number of genes for which the isoform with highest average FPKM was different across centers.

### Supplemental Figures

**Supplementary Figure 1:**
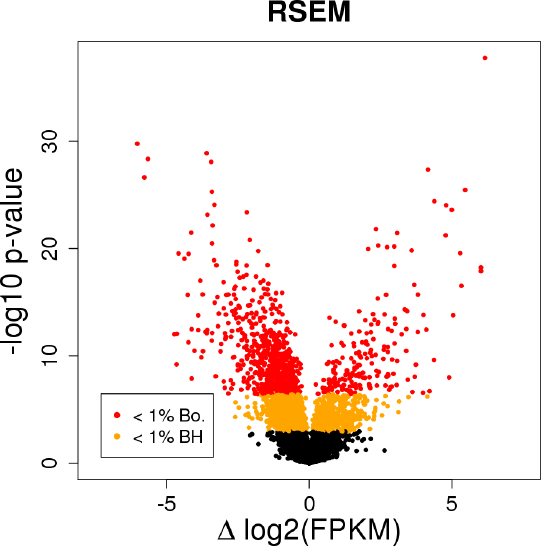
Volcano plot of a comparison of *RSEM* transcript estimates across center. 2,829 transcripts had Benjamini-Hochberg adjusted *p* value less than 1% and 892 had family-wise error rate of 1% using a Bonferroni correction, out of 26,057 transcripts with FPKM estimate greater than 0.1.

**Supplementary Figure 2:**
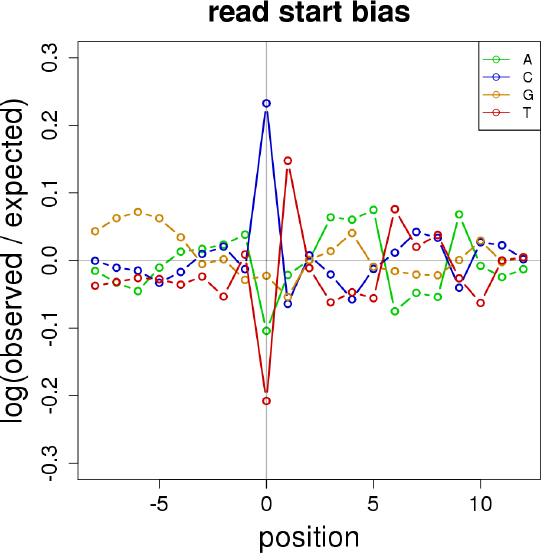
The 0-order terms of the read start bias model estimated for the 5’ fragment end for one sample of the IVT-seq dataset. The 0-order terms are shown for visual simplicity, although the variable length Markov model (VLMM) used here and proposed by Roberts et al. [13] has higher order (1- and 2-order) Markov dependence for the middle positions. Both 5’ and 3’ end biases are combined for the read start bias calculation.

**Supplementary Figure 3:**
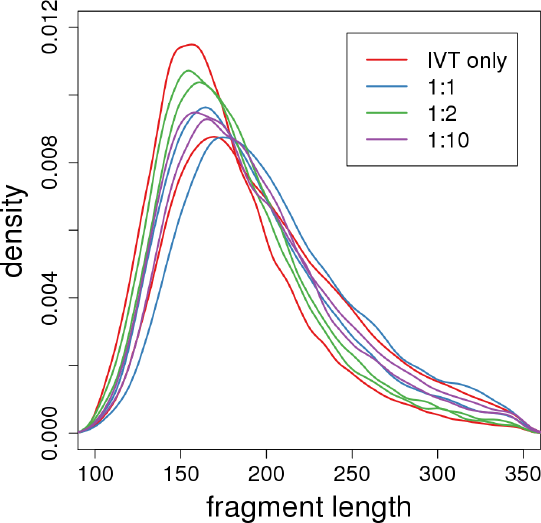
The fragment length distributions for IVT-seq samples.

**Supplementary Figure 4:**
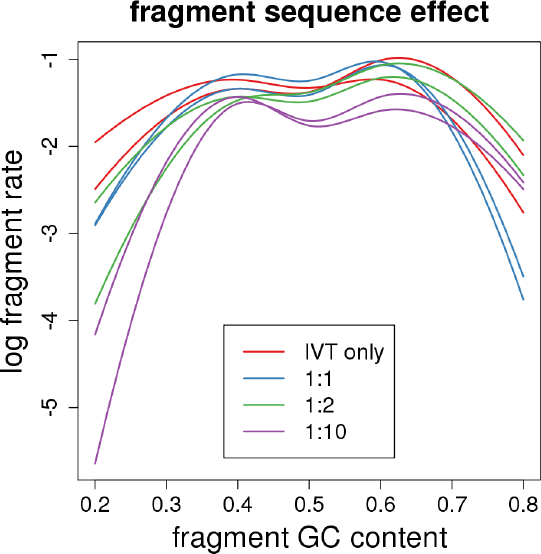
The dependence of fragment rate on fragment GC content for IVT-seq samples. The following smooth curves were fit for the model with all terms, therefore representing the GC content dependence after removing read start bias and fragment length bias (fit as an offset).

**Supplementary Figure 5:**
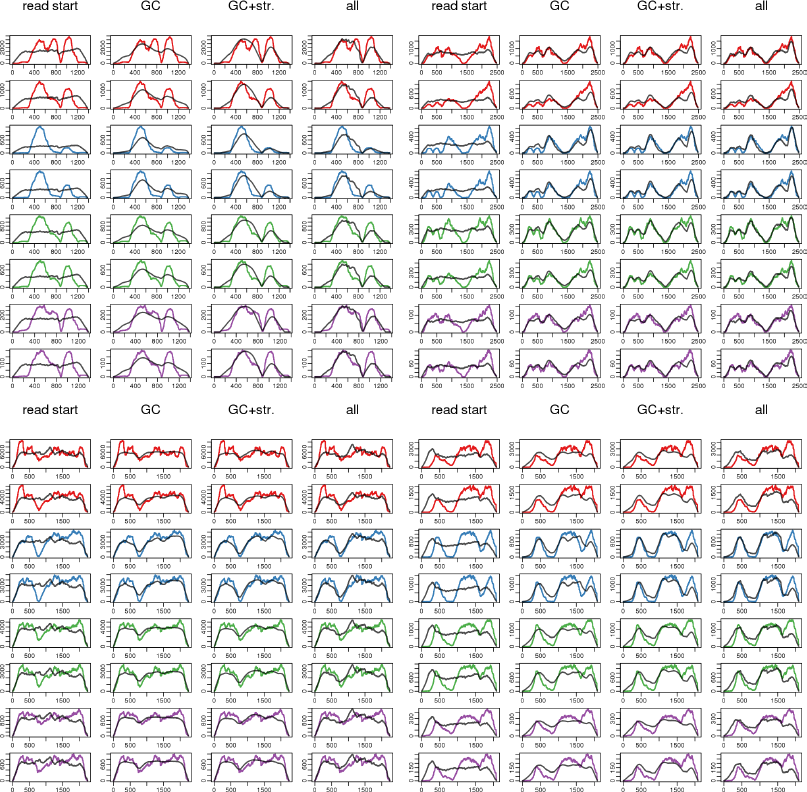
Test set prediction of coverage for IVT-seq transcripts. Predicted coverage (black lines) and raw fragment coverage (colored lines) is shown for four different bias models and four transcripts: BC000158, BC011047, BC011377 and BC011380 (from top left, across rows), and for all eight samples (color denotes sample condition). Test set mean squared error was calculated by averaging the squared residuals from the predicted to the observed coverage.

**Supplementary Figure 6:**
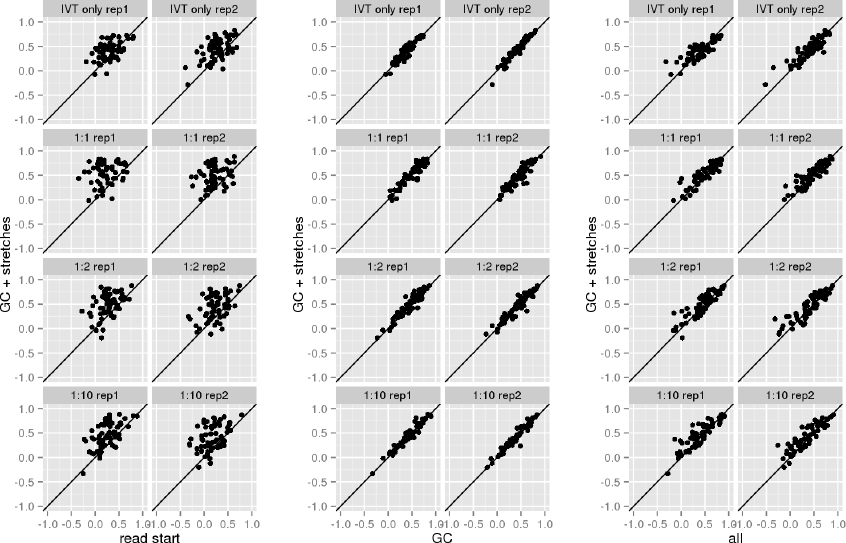
Comparison of the reduction in test set mean squared error (MSE) across the four models, split for eight IVT-seq samples. In all scatterplots, the y-axis shows the reduction in MSE for the model with fragment GC content and GC stretches, compared to a null model of uniform coverage. The x-axis shows (from left to right) the reduction in MSE for the read start model, the fragment GC content model (no GC stretches), and the model with all terms.

**Supplementary Figure 7:**
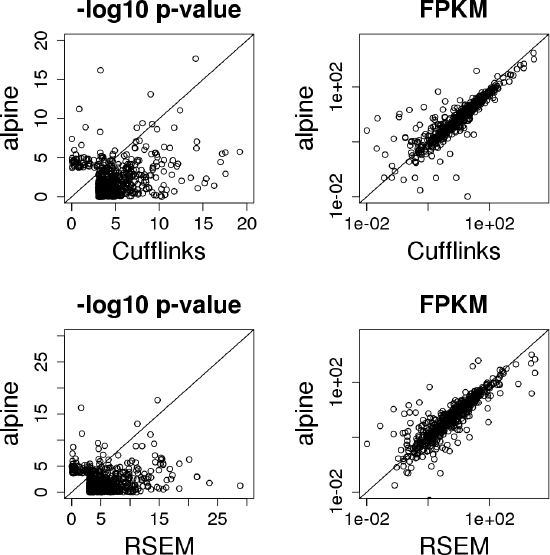
Comparison of *-log*_10_ *p* values and FPKM estimates across methods. Raw *p* values are shown for the set of transcripts with adjusted *p* values less than 0.1. In the top row, *Cufflinks* and *alpine* run with the “GC+str.” bias terms are compared (see Methods). In the bottom row, *RSEM* and *alpine* run with the “GC+str.” bias terms are compared.

**Supplementary Figure 8:**
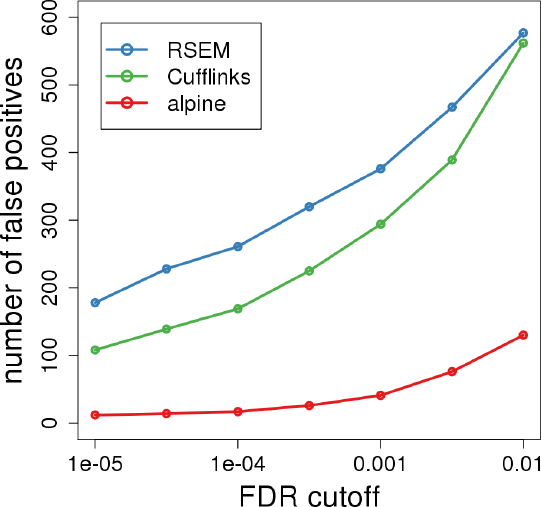
Total number of false positives at various false discovery rate (FDR) cutoffs. Shown are the total number of transcripts reported as differentially expressed among 5,712 transcripts from genes with two isoforms, when comparing log_2_(FPKM+1) estimates of GEUVADIS samples across sequencing center. *p* values were adjusted using the Benjamini-Hochberg method.

**Supplementary Figure 9:**
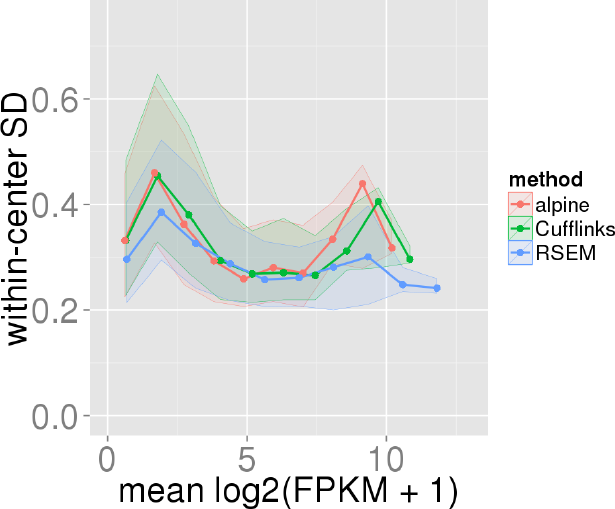
Within-center standard deviation and mean of log_2_(FPKM + 1) estimates. Shown is the median (dark line), and 25% to 75% quantile (shaded region) of within-center standard deviation of transcript estimates for 10 bins along the mean. Only transcripts with FPKM estimates greater than 0.1 are included.

**Supplementary Figure 10:**
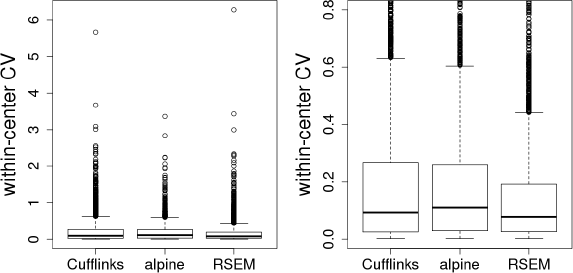
Within-center coefficient of variation of log_2_(FPKM + 1) estimates. For transcripts with FPKM greater than 0.1, the within-center coefficient of variation (variance divided by mean) was calculated for each center and averaged.

**Supplementary Figure 11:**
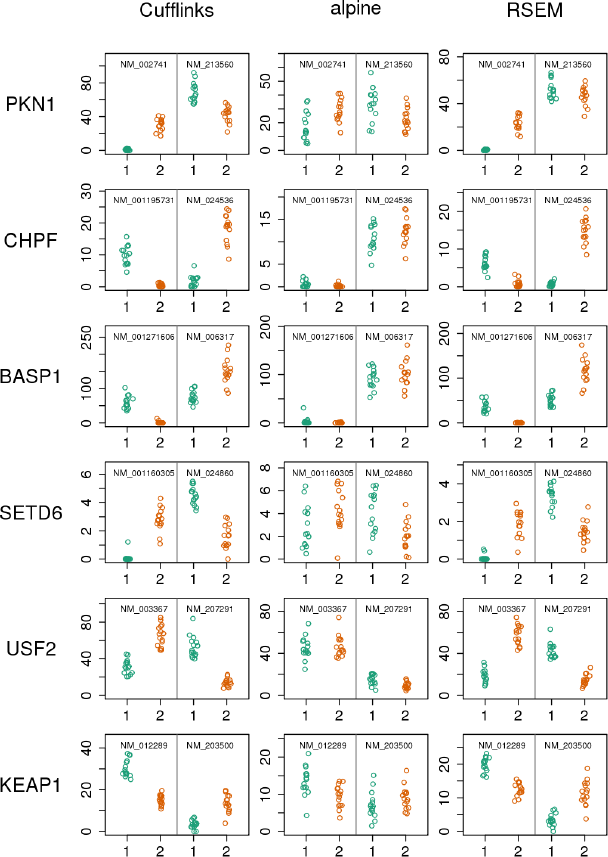
Examples of estimated FPKM across sequencing center for genes with two isoforms, for the three methods. Sequencing center (1 or 2) is indicated on the x-axis. Examples selected for large across-center differences for *Cufflinks* and *RSEM.*

**Supplementary Figure 12:**
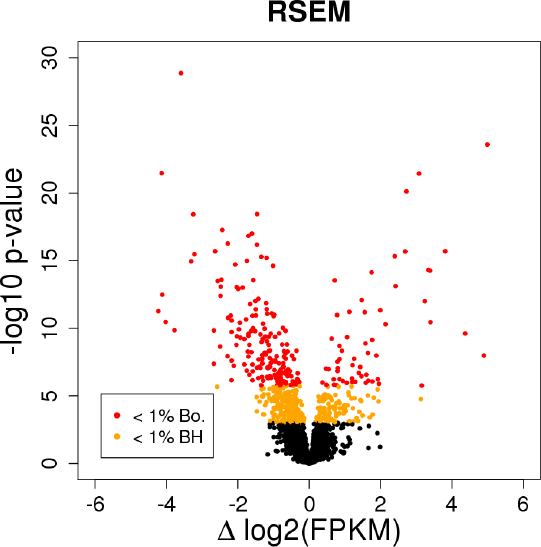
Volcano plots of differential transcript expression across centers for genes with two isoforms, using *RSEM* estimated FPKM. Out of 5,712 transcripts, 577 transcripts reported differential expression across center using an adjusted *p* value threshold of 1%, and 239 transcripts using a conservative Bonferroni family-wise error rate of 1%. 4,979 of the 5,712 transcripts have average FPKM greater than 0.1.

**Supplementary Figure 13:**
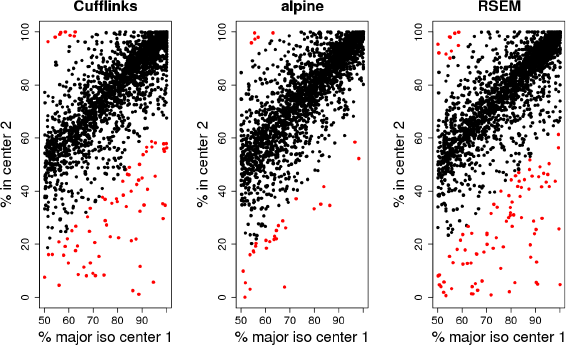
Consistency of percent expression of the major isoform across center across 2856 genes. *Cufflinks* had 78 genes with a change in estimated isoform percent greater than 35%, while *alpine* had 32 genes and *RSEM* had 97 genes.

**Supplementary Figure 14:**
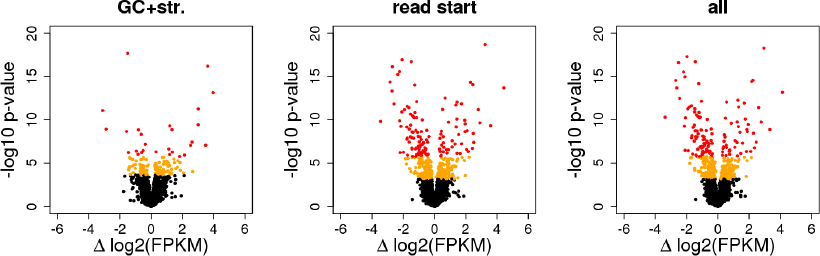
Comparison of across-center differences for different *alpine* bias models. At a threshold on Benjamini-Hochberg adjusted *p* values of 1%, the models reported 130, 393 and 372 transcripts differentially expressed out of 5,712, for the models with GC content + GC stretches, read start, and a model with all terms, respectively. At a more conservative Bonferroni threshold of 1% family-wise error rate, the models reported 27, 125 and 124 transcripts differentially expressed, respectively. See Methods for model details.

**Supplementary Figure 15:**
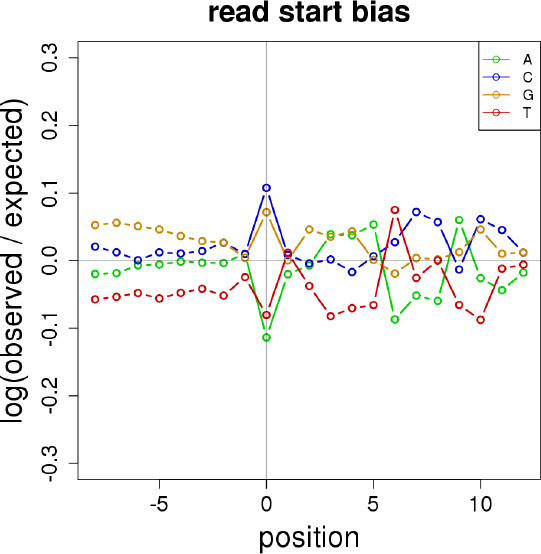
The 0-order terms of the read start bias model estimated for the 5’ fragment end for one sample of the GEUVADIS dataset. As in Supplementary Figure 2, the 0- order terms are shown for visual simplicity, although the variable length Markov model (VLMM) used here has higher order (1- and 2-order) Markov dependence for the middle positions.

**Supplementary Figure 16:**
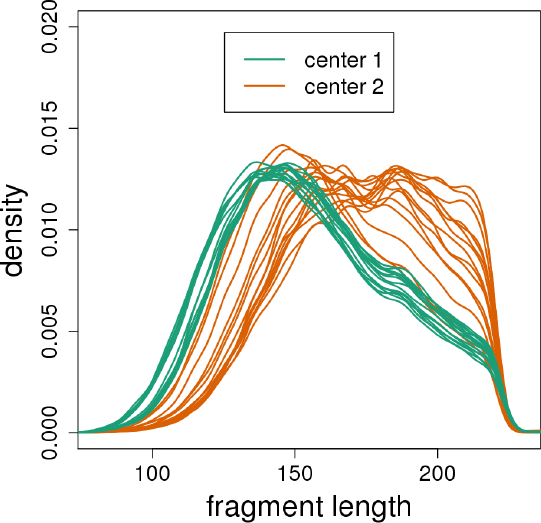
The fragment length densities calculated for GEUVADIS samples.

**Supplementary Figure 17:**
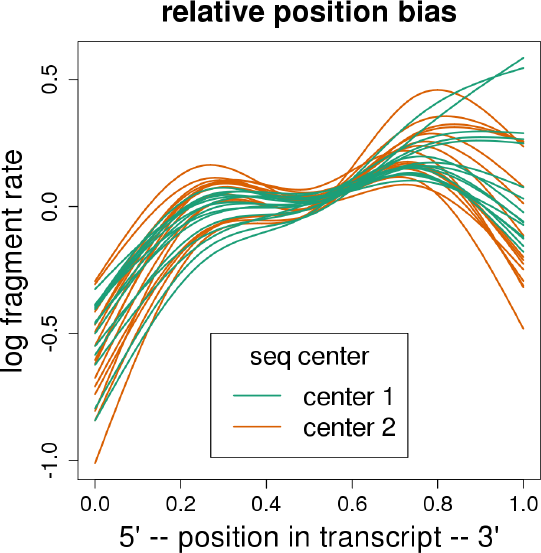
The relative position bias curves calculated for GEUVADIS samples.

